# Evaluation of an H5 influenza virus mRNA-lipid nanoparticle (LNP) vaccine in lactating dairy cows

**DOI:** 10.64898/2026.03.03.709308

**Authors:** Jefferson J. S. Santos, Carine K. Souza, Giovana C. Zanella, Débora B. Goulart, Bailey Arruda, Paola Boggiatto, Mitchell V. Palmer, Celeste A. Snyder, Michaela A. Kristula, Charlene Dickens, Terry L Webb, Reilly K. Atkinson, Bernadeta Dadonaite, Garima Dwivedi, Mohamad-Gabriel Alameh, Jesse D. Bloom, Drew Weissman, Gary C. Althouse, Amy L. Baker, Scott E. Hensley

## Abstract

Highly pathogenic avian influenza (HPAI) clade 2.3.4.4b H5N1 virus has recently emerged in dairy cattle in the United States. The virus replicates primarily in the mammary gland of infected cattle, leading to dramatic reductions in milk production. It is thought that the virus transmits from animal to animal through viral shedding in milk, and therefore, vaccines that decrease the amount of virus in milk can potentially limit the current outbreak and reduce the risk of H5N1 spillover into humans. Here, we assess the immunogenicity and efficacy of a clade 2.3.4.4b H5 mRNA-LNP vaccine in lactating dairy cows. We found that the H5 mRNA-LNP vaccine elicited robust antibody responses in sera and milk and significantly reduced viral replication and disease caused by clade 2.3.4.4b H5N1 intramammary infection.

## Introduction

Highly pathogenic avian influenza (HPAI) H5N1 clade 2.3.4.4b virus continues to cause widespread outbreaks among wild birds and mammals in North America, as well as on commercial poultry farms. H5N1 clade 2.3.4.4b emerged in cattle in the U.S and quickly spread to animals in 19 states^1^. The H5N1 clade 2.3.4.4b outbreak in cattle has already caused major economic losses in the dairy industry ^2^ and represents a concern for human health due to the increased risk of mammalian adaptation and potential human-to-human transmission^3^.

H5N1 clade 2.3.3.4b virus causes limited respiratory symptoms in cattle. While the virus can be sporadically detected from oronasally inoculated calves under experimental conditions^4–6^, the respiratory tract is not likely the primary site of viral replication. Instead, the main site of H5N1 viral replication is the mammary tissue of infected lactating animals. Experimental intramammary inoculation with H5N1 clade 2.3.4.4b viruses leads to reduced feed intake and rumen motility, as well as drastic reductions in milk production due to severe mastitis^4, 5, 7, 8^. H5N1 clade 2.3.4.4b viruses exhibit tropism for the mammary gland tissue of cows^9, 10^, as confirmed by the presence of viral RNA and antigen in the alveolar milk-secreting epithelial cells and the interlobular space of infected animals^8^. Additionally, mastitis caused by the H5N1 clade 2.3.4.4b virus affects the color and consistency of milk, and changes in the mammary tissue contribute to the dramatic reduction in milk production^4^.

In addition to enhanced biosecurity efforts, the use of HPAI clade 2.3.4.4b H5N1 vaccines in cattle has been considered^11^. Vaccination can potentially aid in controlling the current outbreak in lactating dairy cows, limiting the drop in milk production in exposed animals, and reducing the risk of spillover events into humans. Some bovine vaccines elicit antibodies that are transferred into milk in lactating animals. For example, the *M. bovis*-BoHV-1 combined vaccine effectively elicits antigen-specific maternal antibodies^12^, and an inactivated H5 avian influenza A virus vaccine elicits antibodies in both serum and milk^13^. While other vaccines against the HPAI clade 2.3.4.4b H5N1 virus have been tested in bovine animal models^6, 13, 14^, studies are needed to systematically evaluate the immunogenicity and protective efficacy of vaccines in lactating dairy cows against intramammary infection, as the mammary glands are the site for HPAI clade 2.3.4.4b H5N1 virus infection^4, 5^.

We developed an mRNA-LNP vaccine encoding an HA from a clade 2.3.4.4b H5N1 virus and demonstrated that the vaccine elicited robust immune responses in mice and ferrets, preventing morbidity and mortality in ferrets following HPAI 2.3.4.4b H5N1 virus challenge^15^. We found that the same mRNA-LNP vaccine was immunogenic in calves, inducing a strong antibody and T-cell mediated immune response that reduces viral shedding from experimental HPAI 2.3.4.4b H5N1 virus infection^6^. In the present study, we evaluated the immunogenicity and efficacy of a clade 2.3.4.4b H5 mRNA-LNP vaccine administered to lactating dairy cows. We found that the H5 mRNA-LNP vaccine elicited strong antibody responses in both serum and milk. Additionally, the vaccine significantly reduced virus replication and disease, including clinical mastitis, associated with clade 2.3.4.4b H5N1 intramammary route of infection.

## Results

### H5 mRNA-LNP vaccines are well tolerated in lactating dairy cattle

We created a new vaccine that expresses the hemagglutinin (HA) of A/dairy cattle/Texas/24-008749-002/2024, which is representative of HPAI clade 2.3.4.4b H5N1 viruses currently circulating in dairy cattle in the U.S.^4^. We vaccinated 4 lactating dairy cows intramuscularly (i.m.) with 500 μg of H5 mRNA-LNP in saline and then boosted i.m. with the same vaccine dose 3 weeks later (**Fig.1a**). As a control, we i.m. injected 4 additional lactating cows twice with saline (**Fig. 1a**). Animals in our study were obtained from a HPAI-monitored Holstein dairy herd in Pennsylvania and ranged from 2 to 8 years of age (**Supplementary Table 1**). We collected serum samples at 0, 7, 21, 28, 41, and 49 days after the initial vaccination and we collected milk samples daily. Both unvaccinated and vaccinated animals were challenged with infectious clade 2.3.4.4b H5N1 virus through intramammary inoculation 49 days after the first vaccination.

**Figure 1.**
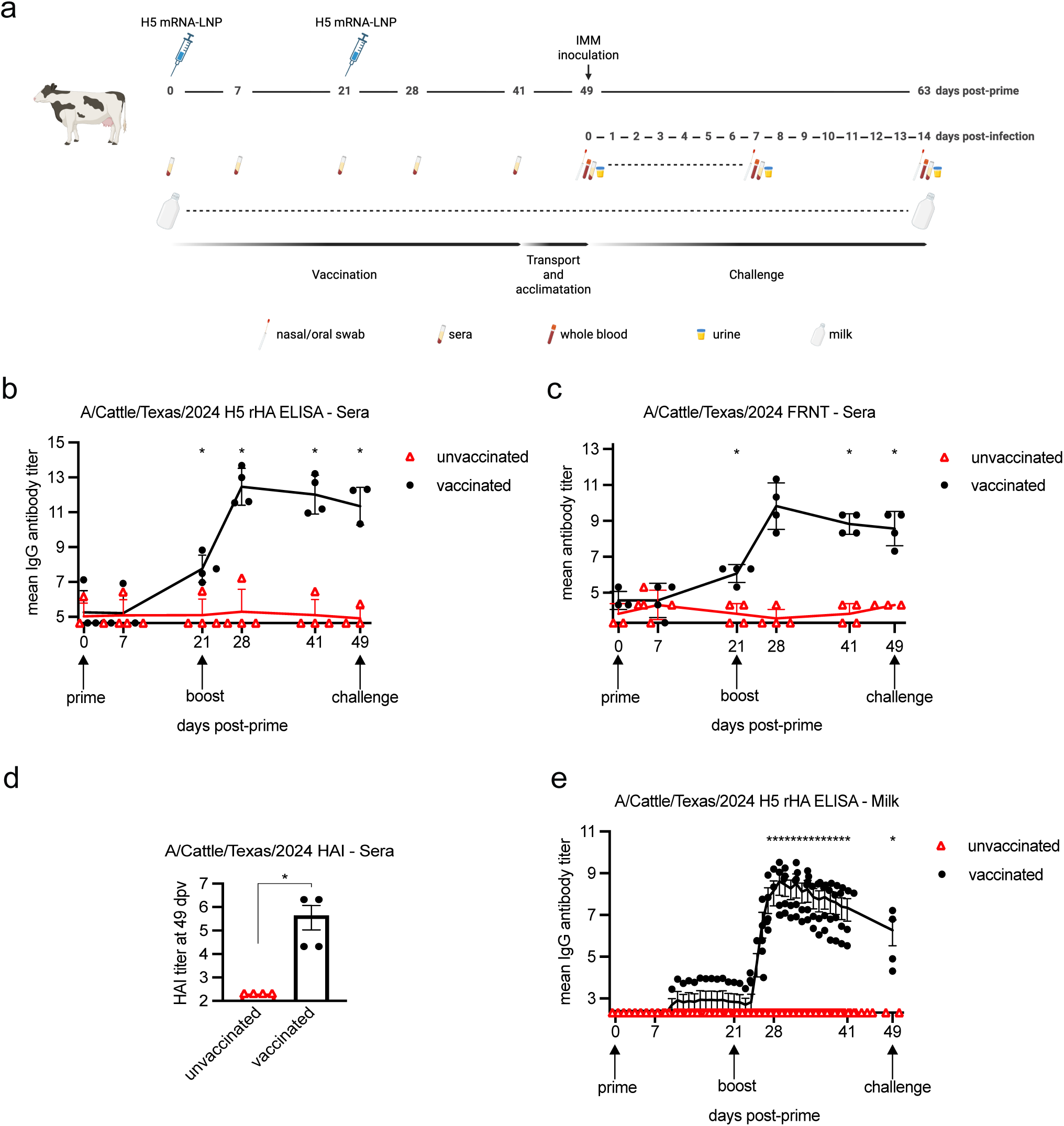
Vaccination study design and mRNA-LNP vaccine immunogenicity in lactating dairy cows. **(a)** Overview of the vaccination and sampling scheme. Lactating dairy cows were vaccinated using a prime-boost regimen. Milk and blood samples were collected at different time points. At 49 dpv and 28 days post-boost, lactating dairy cows were challenged with HPAIV H5N1 and monitored daily for signs of disease and virus shedding. Figure created with BioRender.com. **(b)** H5-specific IgG antibody levels were measured by ELISA in sera from unvaccinated (open red triangles) and vaccinated (closed black circles) animals. **(c)** Neutralizing antibodies were measured using a 90% foci reduction neutralization test (FRNT90); the reciprocal dilutions of serum required to inhibit 90% of virus infection are shown. **(d)** Neutralizing antibodies measured by hemagglutination inhibition (HI) assay. **(e)** H5-specific IgG antibody levels were measured by ELISA in milk. Data are mean ± SEM, and individual data points (n=4) are shown. Significant statistical differences (*P*<0.05) between groups were determined using an unpaired *t*-test and are indicated by an asterisk.

The vaccine was well-tolerated, with no obvious adverse effects. Previous studies found trace amounts of vaccine-related mRNA in human milk in some individuals shortly after SARS-CoV-2 mRNA-LNP vaccination^16–18^. We therefore completed experiments to determine if we could detect H5 mRNA-LNP in bovine milk samples obtained after vaccination. First, we established a sensitive quantitative PCR assay to measure H5 mRNA levels in milk. We spiked a milk sample from an unvaccinated cow with a known concentration of the H5 mRNA-LNP vaccine to assess the sensitivity of our PCR assay (**Extended Data Fig.1a**). We found that the detection limit of our assay was 65 mRNA copies per μL. We then used our quantitative PCR to measure H5 mRNA levels in milk samples collected up to 8 days post-prime or post-boost vaccination. None of the milk samples from vaccinated animals had detectable levels of H5 mRNA (**Extended Data Fig.1b**), demonstrating that vaccine derived mRNA was not present at detectable levels in milk after i.m. vaccination of lactating dairy cows.

### H5 mRNA-LNP vaccine is immunogenic in lactating dairy cattle

We quantified the level of H5-reactive antibodies in the sera from vaccinated animals and performed functional assays to determine if these antibodies could inhibit H5N1 virus infection. We detected low levels of H5-reactive IgG in sera as early as 21 days post-vaccination (dpv) that significantly increased after booster vaccination (**Fig. 1b**). Unexpectedly, we found that one cow from the unvaccinated group and one cow from the vaccinated group had low levels of H5-reactive serum IgG prior to the start of the experiment but was below the limit of detection in the H5 neutralizing assay (day 0, **Fig. 1b-c**). The source herd was under weekly HPAI surveillance prior and during the study and remained HPAI negative, suggesting that the low levels of H5-reactive IgG in these animals was not due to H5N1 infection. A recent study reported that 2.6% of cattle in the U.S. have serological evidence of past exposures to human influenza A virus^19^, and we therefore hypothesized that the low levels of H5-reactive IgG in 2 of the animals in our study were due to cross-reactive antibodies elicited by human influenza virus infections. Consistent with this idea, the 2 animals that had low but detectable H5 IgG antibodies prior to the study also had detectable H1-reactive antibodies (**Extended Data Fig. 2**).

We completed *in vitro* neutralization assays to determine if sera antibodies could prevent infection of the A/dairy cattle/Texas/2024 H5N1 virus. Consistent with the IgG binding data, H5N1 neutralizing antibodies were detected as early as 21 dpv and were significantly increased after booster vaccination (**Fig. 1c**). We also completed hemagglutination-inhibition (HAI) assays using sera collected just prior to viral challenge to determine if vaccine-elicited antibodies blocked viral attachment. Consistent with both the IgG binding and *in vitro* neutralization data, we found that antibodies in the sera from all 4 vaccinated animals blocked viral attachment (**Fig. 1d**).

Finally, we quantified H5-reactive IgG levels in milk samples collected in our study. H5-reactive IgG was detected in only one vaccinated cow after the 1^st^ vaccination; however, we detected high levels of H5-reactive IgG in milk from all 4 vaccinated animals following the booster vaccination (**Fig. 1e**). Peak levels of H5-reactive IgG were higher in sera compared to milk, and antibody levels declined more rapidly in milk compared to the sera (**Extended Data Fig. 3**). We were unable to perform *in vitro* neutralization with milk samples since non-specific inhibitors in milk from unvaccinated animals inhibit virus in these assays. However, the IgG levels in the milk (**Fig. 1e**) were similar to sera IgG levels (**Fig. 1b**) that were sufficient for neutralization (**Fig. 1c**).

### Specificity of antibodies elicited by H5 mRNA-LNP vaccination

We performed pseudovirus deep mutational scanning experiments^20^ to identify HA epitopes targeted by neutralizing antibodies in sera from lactating dairy cows vaccinated with the H5 mRNA-LNP vaccine (**Fig 2**). For these experiments, we incubated bovine sera with a pool of non-replicative lentiviral particles that were pseudotyped with mutant HAs from the clade 2.3.4.4b A/American Wigeon/South Carolina/USDA-000345-001/2021 virus, as previously described^21^. This H5 deep mutational scanning library carries nearly all of the 10,773 possible amino acid mutations in the HA^21^. We found that the neutralizing activity of the sera from all 4 vaccinated animals targeted the antigenic sites on the globular head of HA (**Fig. 2a-b**), including antigenic site A (residues 124-127 and 144-145, H3 numbering) and antigenic site B (residues 165, 167, 169, and 171, H3 numbering). There was a remarkable similarity in the amino acid sites targeted by the serum antibodies isolated from individual vaccinated cows. These data indicate that most neutralizing antibodies elicited by the H5 mRNA-LNP vaccine target neutralizing epitopes adjacent to the receptor binding domain of HA. Importantly, the amino acid identities at the sites most strongly targeted by the vaccine-induced neutralizing antibodies have not changed substantially in the HA of circulating clade 2.3.4.4b viruses (**Fig. 2c**).

**Figure 2.**
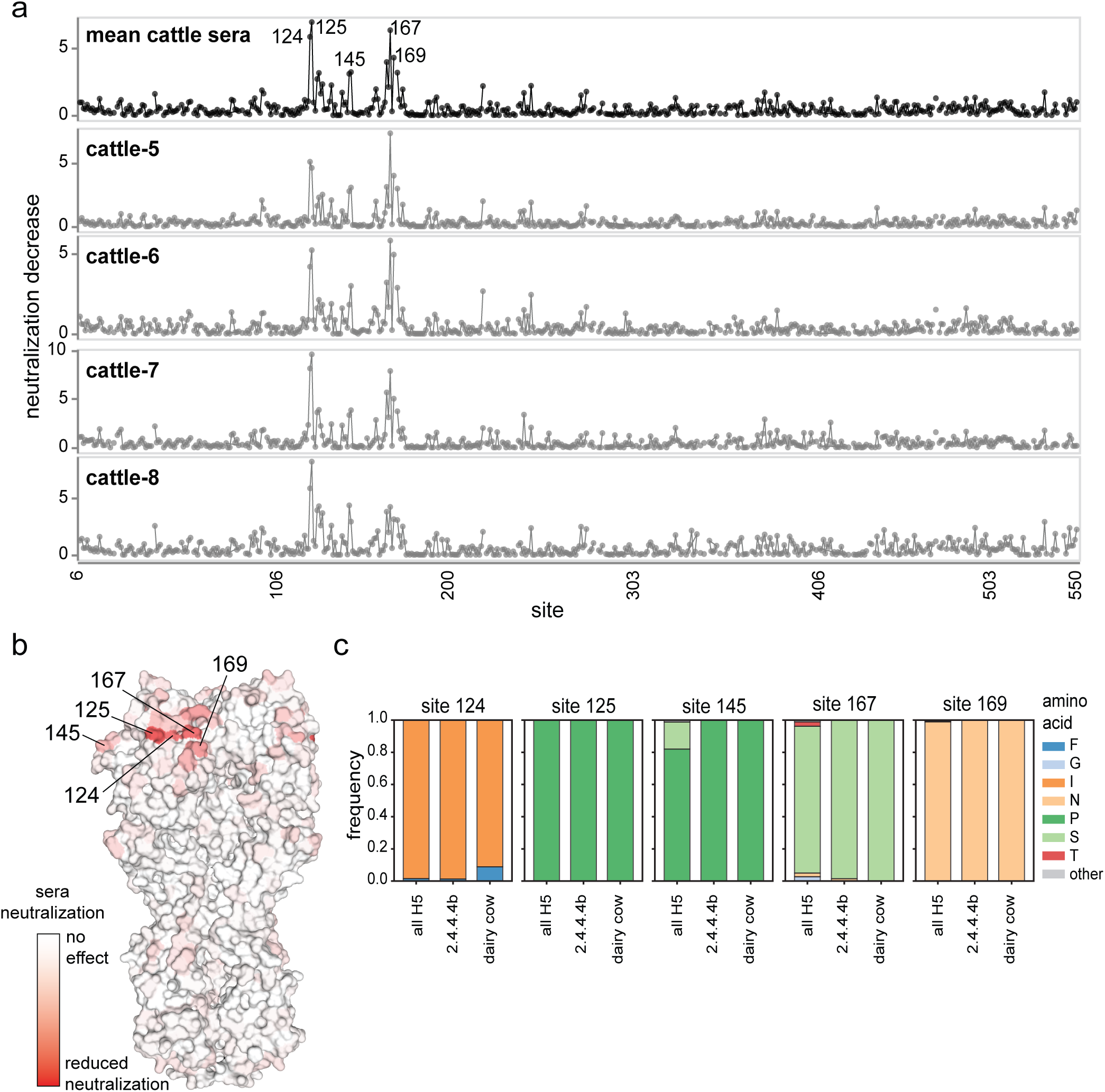
Effect of mutations to HA on vaccinated cattle sera neutralization. **(a)** Neutralization decrease caused by mutations at each position in HA. The top plot shows the mean total neutralization decrease for the four vaccinated cow sera, and the plots below show the individual cow neutralization decrease. Sites causing the greatest neutralization decrease are labeled by amino acid position above the top plot. See https://dms-vep.org/Flu_H5N1_American-Wigeon_2021_HA_DMS_cattle_Penn_sera/htmls/cattle_sera_escape_faceted.html for an interactive version of this plot. **(b)** Mean effects of mutations at each site cattle sera neutralization overlaid on a surface representation of HA (PBD: 4KWM). Sites are colored white if mutations have no effect on neutralization, and red if mutations decrease neutralization. **(c)** Amino acid conservation at HA sites where mutations caused the greatest neutralization decrease by the vaccinated cow sera, calculated over all available H5 sequences, 2.3.4.4b clade sequences, and B3.13 and D1.1 genotype sequences from dairy cows. This figure uses H3 site numbering in the mature HA peptide (see https://dms-vep.org/Flu_H5_American-Wigeon_South-Carolina_2021-H5N1_DMS/numbering.html).

### H5 mRNA-LNP vaccination protects lactating cows from H5N1 infection

Lactating cows infected with HPAI clade 2.3.4.4b H5N1 display virus-induced mastitis, reduced feed intake and rumen motility, decreased milk production, abnormal milk color and consistency, and virus shedding in milk^4, 7, 8^. To determine whether H5 mRNA-LNP vaccination can prevent virus shedding and reduce H5N1 disease, we experimentally challenged unvaccinated and vaccinated cows with clade 2.3.4.4b H5N1 (A/dairy cattle/Texas/24-008749-002/2024) via the intramammary inoculation route and monitored the cows for 14 days post-inoculation (dpi). We then euthanized the animals and performed macroscopic and microscopic lesion analyses.

We measured levels of H5N1 viral RNA in milk samples by quantitative RT-PCR (RT-qPCR) assays. In unvaccinated animals, H5N1 viral RNA in milk from inoculated mammary quarters was first detected at 1 dpi, peaked at 2 dpi, and milk samples remained RT-qPCR-positive throughout the study period (**Fig. 3a**). In vaccinated animals, H5N1 viral RNA levels in milk from inoculated mammary quarters were significantly lower at 1-3, 5, and 8-13 dpi, with ∼1,000-fold lower peak titer compared to levels found in milk from unvaccinated animals (**Fig. 3a**). Most milk samples that were positive for H5N1 viral RNA by RT-qPCR with Ct<35 contained infectious H5N1 virus (**Supplementary Table 2**). We detected either H5N1 viral RNA or infectious virus in milk from the uninoculated quarters of 2 unvaccinated animals (**Extended Data Fig. 4, Supplementary Table 2**), suggesting that virus was able to migrate from an inoculated to uninoculated quarter in these animals. We detected viral antigen in mammary tissue from uninoculated quarters of both unvaccinated and vaccinated animals via immunohistochemistry (IHC) (**Extended Data Fig. 4 and 5a-b**); however, we did not detect viral RNA in the milk from uninoculated mammary quarters in vaccinated animals, suggesting that the vaccine prevented further spread of the virus.

**Figure 3.**
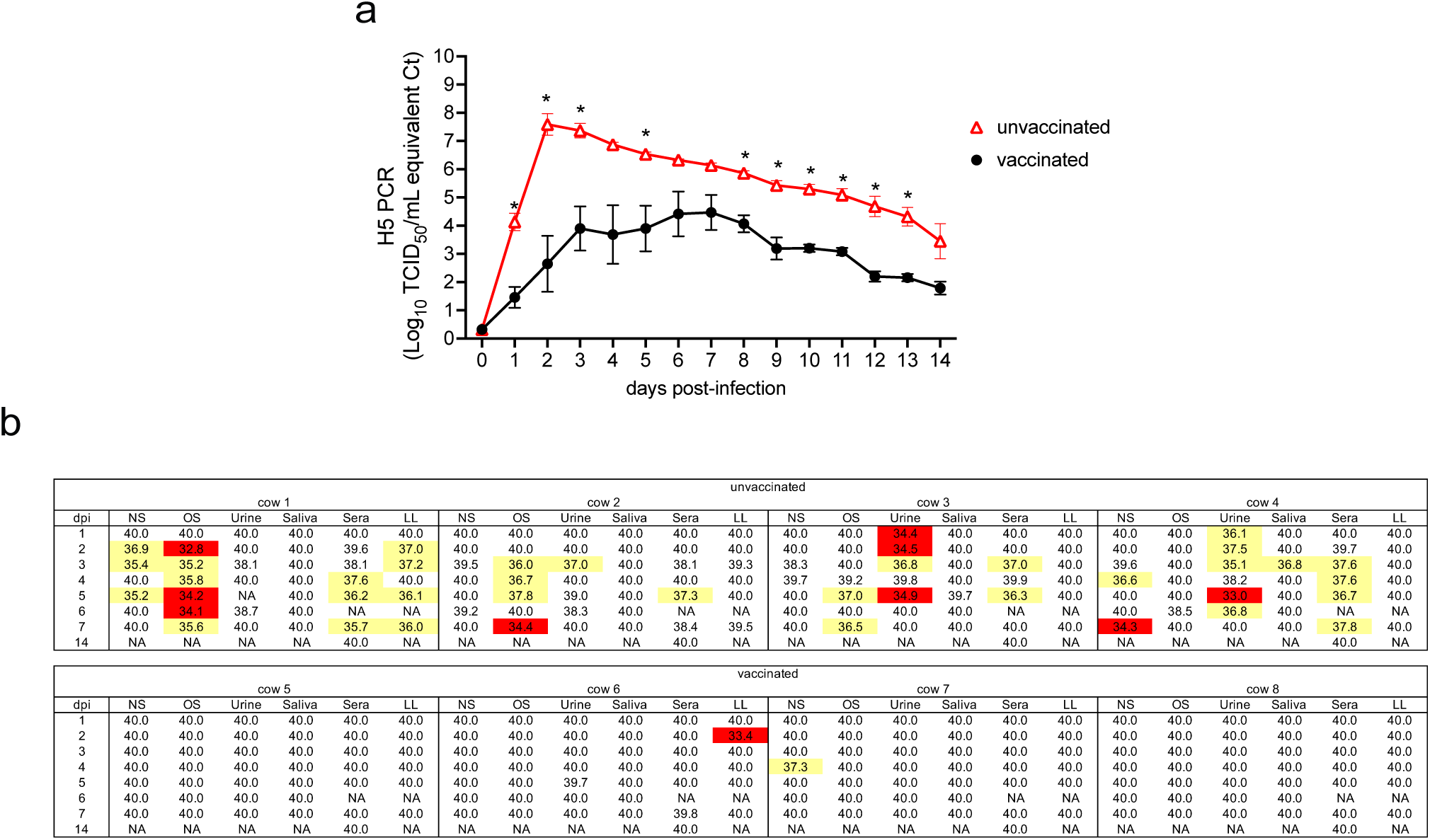
Virus RNA detection in clinical samples from unvaccinated and vaccinated lactating cows after challenge. **(a)** Log_10_ TCID_50_/mL equivalent Ct detection of virus RNA in stripped milk from inoculated mammary quarters of unvaccinated (open red triangles) and vaccinated (closed black circles) lactating dairy cows. Data shown are mean ± SEM of n=4. Time points with significant difference (p<0.05) between unvaccinated and vaccinated indicated by an asterisk. **(b)** Real-time qPCR quantification of virus RNA from antemortem clinical samples of unvaccinated and vaccinated lactating dairy cows. A Ct below 35 was considered positive (red), between 35 and 38 suspect (yellow), and above 38 negative (white). NS, nasal swab; OS, ocular swab; LL, LeucoLOCK from whole blood; NA, not applicable; dpi, days post-inoculation.

We measured H5N1 RNA levels in other clinical samples, including nasal swabs, ocular swabs, urine, saliva, serum, and whole blood. We detected H5 RNA in multiple sample types from all 4 unvaccinated animals (**Fig. 3b, Extended Data Fig 4**). In contrast, few samples obtained from vaccinated animals possessed detectable levels of H5 RNA (**Fig. 3b, Extended Data Fig 4**). We also collected fresh mammary tissue, supramammary lymph nodes, and a section of diaphragm at the 14 dpi necropsy. H5 RNA was detected in mammary tissue from inoculated quarters in all unvaccinated cows, in an uninoculated quarter of one unvaccinated cow, and only in inoculated quarters of 2 vaccinated cows (**Extended Data Fig 4**). Supramammary lymph node tissue was positive for H5 RNA in 3 of unvaccinated cows and in 2 vaccinated cows and diaphragm liquid was suspect in one unvaccinated cow (**Extended Data Fig 4**).

Following H5N1 virus infection, unvaccinated cows displayed decreased rumen motility between 1-7 dpi, with a significant decrease at 3 and 4 dpi (**Fig. 4a**). Milk production also significantly decreased in unvaccinated animals following H5N1 infection, beginning at 2 dpi, with a dramatic drop at 3 dpi that remained significantly reduced throughout the study at 4-8 and 10-12 dpi (**Fig. 4b**). Rumen motility and milk production was less impacted in vaccinated cows following H5N1 infection (**Fig. 4a-b**). We observed severe changes in color (**Fig. 4c**) and consistency (**Fig. 4d**) indicative of mastitis in milk from the inoculated mammary quarters of unvaccinated cows following H5N1 infection. In contrast, milk color and consistency from inoculated mammary quarters of vaccinated cows were less affected by H5N1 infection (**Fig. 4c-d**).

**Figure 4.**
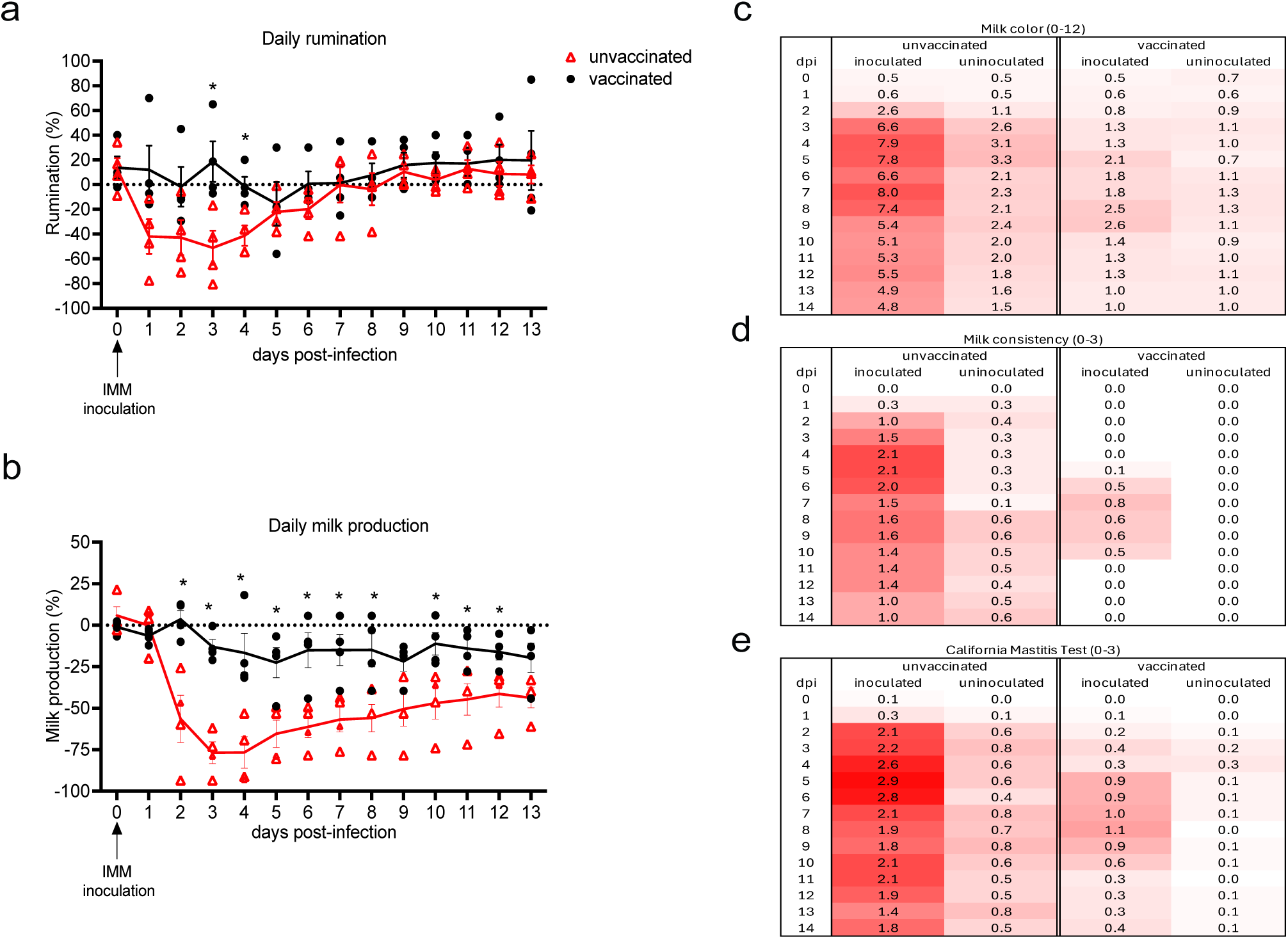
Clinical signs in unvaccinated and vaccinated lactating dairy cows after challenge. Unvaccinated (open red triangles) and vaccinated (closed black circles) lactating dairy cows were challenged with HPAI clade 2.3.4.4b H5N1 and monitored daily for **(a)** rumination and **(b)** milk production. Data are mean ± SEM, and individual data points (n=4) are shown; asterisks indicate significant difference (p<0.05). Collected milk samples were evaluated by quarter for mastitis by **(c)** milk color, **(d)** milk consistency, and **(e)** California Mastitis Test (CMT). Darker red shades represent higher scores; lighter to white shades represent lower or zero scores. Data shown are mean of n=4 per treatment group.

Uninoculated mammary quarters of unvaccinated cows also displayed changes in milk color and consistency, although to a lesser extent compared to inoculated mammary quarters (**Fig. 4c-d**). Further, unvaccinated cows showed signs of mastitis in inoculated quarters, and to a lesser extent in uninoculated quarters, as determined by California Mastitis Test (CMT) (**Fig. 4e**). Inoculated quarters from unvaccinated animals consistently scored above 2 (score range 0-3) in the CMT, which indicates moderate to severe inflammation and leukocyte infiltration. Conversely, inoculated and uninoculated mammary quarters of vaccinated cows frequently scored a CMT below 1 following H5N1 virus infection (**Fig. 4e**), indicating little to no signs of mastitis. The clinical mastitis scores in uninoculated quarters were primarily contributed by 2 unvaccinated cows (**Fig. 4 c-e**). We found that unvaccinated cows had a significant association between loss in milk production clinical signs, viral RNA detections in milk of inoculated quarters. In contrast, vaccinated cows did not present significant association between loss in milk production, clinical signs and viral RNA in milk, except for rumination time (**Extended Data Fig. 6a-b**).

### H5 mRNA-LNP vaccination of lactating cows significantly decreased mammary fibrosis

We completed macroscopic evaluation of the mammary glands from inoculated and uninoculated quarters for all lactating dairy cows at necropsy. Consistent with the clinical observations, inoculated mammary quarters from unvaccinated cows exhibited significantly higher macroscopic lesion scores compared to vaccinated cows (*P<0.05*, 95.4 and 8.1 respectively; **Fig. 5a**). In contrast, macroscopic lesion scores in uninoculated mammary quarters were low and did not differ significantly between unvaccinated and vaccinated cows (*P>0.05*, 12.1 to 10.0; **Supplementary Table 3**). A representative image of normal mammary tissue in an uninoculated quarter is shown in **Fig. 5b** and an image of fibrotic mammary tissue in an inoculated quarter of an unvaccinated cow is shown in **Fig. 5c**. Consistent with the macroscopic scores, microscopic scores of mammary gland fibrosis using Picrosirius Red staining demonstrated less fibrosis in inoculated glands from vaccinated animals compared to unvaccinated animals (**Fig. 5d-h**). A histologically unremarkable photomicrograph of mammary tissue is shown in **Fig. 5e** for comparison, and representative photomicrographs are shown with tissue from a vaccinated uninoculated cow (**Fig. 5f**), a vaccinated inoculated cow (**Fig. 5g**), and an unvaccinated inoculated cow (**Fig. 5h**).

**Figure 5.**
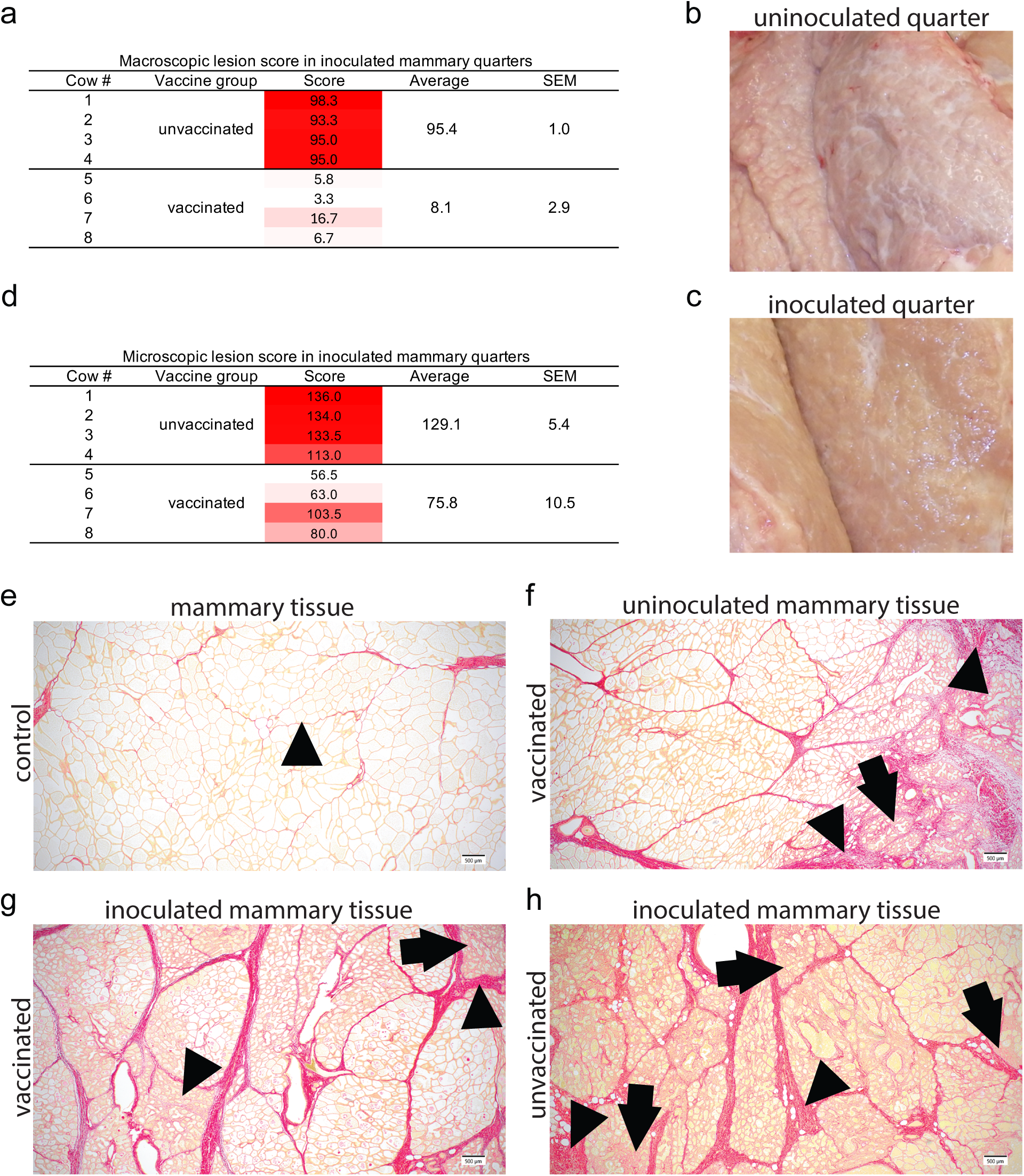
Macroscopic and microscopic lesions in mammary tissue after challenge. **(a)** The percentages of macroscopic lesions in the mammary tissue of unvaccinated and vaccinated lactating dairy cows were measured in inoculated quarters. Data shown is the calculated average percentage (0-100) of the rostral, mid, and caudal regions for inoculated quarters, mean ± SEM of n=4. **(b)** Macroscopic photo representative of a normal mammary tissue producing milk. **(c)** Representative photo of mastitis-affected mammary tissue with fibrosis (Cow 2, (b) unvaccinated rear right uninoculated quarter, and (c) rear left inoculated quarter). **(d)** The percentage of microscopic lesions in the mammary tissue of unvaccinated and vaccinated lactating dairy cows. Data shown is the calculated average score of the rostral, mid, and caudal regions for inoculated quarters (scale 0-144), mean ± SEM of n=4. **(e)** Picrosirius Red stain of mammary gland highlighting connective tissue and fibrosis (red). A histologically unremarkable photomicrograph was included for comparative purposes. Thin bands of supporting connective tissue (arrowhead) separate lobules and no to minimal intralobular collagen (Cow 6, front left quarter, vaccinated, uninoculated, fibrosis Score 0). Representative photomicrographs were selected based on their alignment with the mean fibrosis score for the respective group. **(f)** Extensive interlobular fibrosis with lobular separation (arrowhead) and dense intralobular collagen (arrow) with partial lobular disruption (Cow 5, front left quarter, vaccinated uninoculated, fibrosis score 9). **(g)** Thick interlobular collagen bundles with compression of adjacent lobules (arrowhead) and increased intralobular collagen (arrow) with mild distortion of lobular architecture (Cow 7, front right quarter, vaccinated inoculated, fibrosis score 9). **(h)** Extensive interlobular fibrosis with lobular separation (arrowhead) and dense intralobular collagen (arrow) with near complete lobular disruption (Cow 4, rear left quarter, unvaccinated inoculated, fibrosis Score 14).

## Discussion

The recent widespread outbreaks of HPAI clade 2.3.4.4b virus in dairy cattle, along with its association with milk loss and severe mastitis, require effective countermeasures for the dairy industry. In one study, decreased milk production, mortality, and early herd removal of lactating cows clinically affected by HPAI clade 2.3.4.4b virus resulted in economic losses of $950 per clinically affected cow, totaling approximately $737,500 for the infected herd that was studied^2^. Vaccines that reduce clade 2.3.4.4b H5N1 virus in milk might be useful in preventing the spread of clade 2.3.4.4b H5N1 viruses in lactating cattle. Combined with robust viral testing programs, an 2.3.4.4b H5N1 vaccine has the potential to reduce clinical signs and mastitis in H5N1-infected lactating dairy cattle, while limiting potential viral spillovers into the human and other susceptible wild and domestic animal populations.

Our study provides a comprehensive assessment of an H5 mRNA-LNP vaccine against intramammary HPAI clade 2.3.4.4b H5N1 virus infection in lactating dairy cows. We found that our H5 mRNA-LNP vaccine induced neutralizing antibodies in lactating dairy cows and protected against intramammary HPAI clade 2.3.4.4b H5N1 virus infection. The vaccine was well-tolerated, and animals showed no apparent adverse events following vaccination. We did not detect H5 mRNA in milk collected after intramuscular vaccine administration, demonstrating its safety for cows and their milk. The H5 mRNA-LNP vaccine elicited robust antibody responses in both serum and milk. Antibody levels in milk were consistently lower compared to those in serum, prior to challenge. This was expected, considering the varying concentrations of IgG present in milk after the periparturient and colostrogenesis periods. Future studies are needed to investigate the impact of different routes of immunization, vaccination schedules, and vaccine doses on the magnitude and longevity of antibody responses elicited by mRNA-LNP vaccines in cattle. It is critical to continue optimizing vaccine regimens that elicit sustained protective levels of antibodies in the milk of lactating cattle. Additionally, it will be important to consider a differentiating infected from vaccinated animals (DIVA) strategy for diagnostics and HPAI outbreak investigation in vaccinated livestock. Since our H5 mRNA-LNP vaccine elicited only antibody responses against the HA protein, the detection of NP antibodies would indicate a natural infection and could be used as a DIVA strategy.

HPAI H5N1 2.3.4.4b has a high tropism for the mammary gland and infects milk-secreting mammary epithelial cells lining the alveoli resulting in a viral-induced mastitis^8^. Experimental inoculation of lactating cows with HPAI H5 2.3.4.4b via the intramammary route recapitulates field observations, leading to severe acute mammary gland infections with necrotizing mastitis^5^. In affected mammary tissue, 15% to 50% of the secretory alveoli and ducts, both intralobular and interlobular, are replaced by fibrous connective tissue^4^. In our studies, we used a special collagen stain to directly compare fibrosis in mammary tissue caused by H5N1 infection. The vaccinated cows developed significantly less mammary fibrosis in inoculated quarters compared to unvaccinated cows. Although we were unable to confirm a prior history of mastitis for the cows used in this study, mild fibrotic changes to the mammary gland of dairy cattle are not unexpected nor do they severely affect milk production. Vaccinated cows showed mild mammary fibrosis that could be due to a previous mastitis episode or to our H5N1 2.3.4.4b virus inoculation, but this did not affect milk production as compared to the unvaccinated cows. Our work highlights that the H5 mRNA-LNP 2.3.4.4b vaccine provided protection against severe fibrosing mastitis induced by H5N1 infection that will likely protect milk yields across subsequent lactations.

Previous experimental challenge studies reported that the 2.3.4.4b H5N1 genotype B3.13 virus did not migrate to neighboring uninoculated mammary glands^4, 5, 22^. However, in our study, we found evidence that virus migrated from an inoculated to uninoculated mammary quarters of unvaccinated cows following 2.3.4.4b H5N1 infection. Viral RNA was detected in milk samples and mammary gland tissue, virus was isolated from milk samples, and viral antigen was detected by IHC in mammary gland tissues of uninoculated quarters. Additionally, we observed mild clinical signs of mastitis in corresponding uninoculated quarters of unvaccinated cows. Although we also detected viral antigen by IHC in an uninoculated quarter of one vaccinated cow, we did not detect evidence of virus or viral RNA in milk or in any other sample type from this animal. These data suggest that vaccination prevented further spread of virus within the uninoculated quarter, and furthermore, prevented spread of virus outside of the mammary gland and draining lymph nodes. Our study is the first to report 2.3.4.4b H5N1 genotype B3.13 virus migration between mammary quarters, and we found that vaccination reduced viral migration in our studies. Additional studies should evaluate if other H5N1 vaccine candidates prevent the migration of 2.3.4.4b H5N1 to neighboring uninoculated mammary glands.

It will be important to monitor future antigenic changes in clade 2.3.4.4b viruses in cattle. Most of the sera neutralizing antibodies elicited by H5 mRNA-LNP vaccination in our study targeted antigenic sites A and B in the globular head of HA near the receptor binding site. These sites have remained antigenically stable within clade 2.3.4.4.b viruses, and continued surveillance studies will be useful to identify any future changes in these sites. While we did not assess T cell-mediated immunity in the current study, we previously found that mRNA-LNP vaccination induces a potent CD8+ T cell response in calves^6^. Thus, our H5 mRNA-LNP vaccine is well-matched to currently circulating clade 2.3.4.4.b viruses and also elicited T cell responses that may be protective if these viruses evolve to escape antibody responses in the future.

In conclusion, our study demonstrates that the H5 mRNA-LNP 2.3.4.4b vaccine is safe and immunogenic in lactating dairy cows, eliciting robust antibody responses in sera and milk. Moreover, the H5 mRNA-LNP vaccine reduced clinical disease, particularly milk loss, and significantly reduced virus shedding in milk and mammary gland fibrosis in vaccinated lactating cows. The H5 mRNA-LNP 2.3.4.4b vaccine in dairy cows provides an additional tool to use in combination with current control measures to limit the outbreak on farms, reduce spread between farms, and alleviate the economic losses to the dairy industry.

## Online Methods

### Ethical compliance

The Institutional Animal Care and Use Committees of the University of Pennsylvania Office of Animal Welfare and USDA-Agricultural Research Service (ARS) National Animal Disease Center (NADC) reviewed and approved all studies.

### Study Design

Eight Holstein lactating cows at 2-8 years of age (**Supplementary Table 1**) were split into two groups at the New Bolton Center Marshak dairy herd at the University of Pennsylvania and received either saline (unvaccinated) or 500 μg of A/dairy cattle/Texas/24-008749-002/2024 (H5N1) HA mRNA-LNP in saline vaccine (vaccinated) intramuscularly, directly into the muscle of the neck in front of the shoulder and below the nuchal ligament. The vaccine regimen consisted of two doses of the mRNA-LNP vaccine administered with a three-week interval between doses. The vaccine was created using standard techniques as previously described^15^. Blood samples were collected at 0, 7, 21, 28, 41, and 49 dpv, while milk samples were collected daily. Vaccinated cows were milked segregated away from the herd and their milk did not enter the human food supply. At 42 dpv, cows were transported from the New Bolton Center at the University of Pennsylvania to the USDA-ARS NADC in Ames, IA. At 49 dpv, cows were inoculated with 1 ml of 1 x 10^5^ tissue culture infectious dose 50 (TCID_50_) per ml of A/dairy cattle/Texas/24-008749-002/2024 (H5N1) via the intramammary route in each of two quarters, using a teat canula^4^. The inoculation occurred after milking and standard teat disinfection, and the teats were additionally disinfected with an alcohol wipe immediately before the procedure. After infusion, the inoculum was manually moved upwards into the teat sinus with gentle massage. Antemortem samples such as nasal swab, ocular swab, urine, saliva, blood serum (BD vacutainer SST), and blood in EDTA (BD EDTA K2) were collected from 0 to 7 days post-inoculation (dpi). Milk samples from each quarter were collected daily up to 14 dpi. Lactating cows were monitored continuously for rumination using an ear-tag accelerometer sensor from -6 to 14 dpi (Cow Manager SensOor, Agis Automatisering). Cows were recorded daily for rectal temperature and visually monitored for clinical signs, including cough, nasal and ocular discharge, and fecal consistency. Milk samples were collected by quarter and evaluated for mastitis by the California mastitis test (CMT, ImmuCell), for consistency, and for color using a visual scorecard on a scale of 0–12^4^. Samples were also collected from the bucket following milking. At 14 dpi, cows were sedated using an alpha 2-adrenergic receptor antagonist, xylazine (Rompun, Dechra), at 0.1 mg/kg, administered intramuscularly (IM). Animals were then anesthetized using ketamine (Zetamine, VetOne), at 2 mg/kg administered intravenously (IV), and humanely euthanized via IV pentobarbital (Fatal-Plus, Vortech) administration per manufacturer’s dosage recommendation. At necropsy, mammary tissue from each quarter, supramammary lymph nodes, and diaphragm were collected for pathologic evaluation and virus detection by PCR as described below. The study was carried out in a working dairy farm facility during the vaccination stage and in an Agricultural Biosafety level 3 (BSL3-Ag) facility during the challenge stage.

### Vaccine mRNA undetected in milk

After prime and boost immunizations, milk samples were collected and tested for the presence of residual H5 mRNA from the vaccine. RNA was isolated from 50ul of milk using the MagMAX-96 viral RNA isolation kit (Thermo Fisher Scientific). The RT-qPCR assay was performed in a QuantStudio 3 real-time PCR instrument (Thermo Fisher Scientific) using the TaqMan fast virus 1-step master mix for qPCR (Thermo Fisher Scientific) following the manufacturer’s instructions and the primers/probe designed with the PrimerQuest Tool (IDT). The position and sequence of primers and probe used for the assay are reported in **Supplementary Table 5**. A standard curve was generated by spiking a milk sample with a known concentration of the H5 mRNA-LNP vaccine. A 10-fold serial dilution curve, ranging from 8 × 10^-0.1^ to 10^10.9^ mRNA copies μl^-1^ of template, was used to correlate cycle threshold (Ct) values with the amount of mRNA in milk. The limit of detection of the assay was assessed as 8 × 10^0.9^ mRNA copies μl^-1^ of template (65 mRNA copies per μl). All samples and standards were tested in duplicate.

### Recombinant HA (rHA) protein expression

The extracellular domain of A/dairy cattle/Texas/24-008749-002-v/2024(H5N1) HA was synthetically fused to the trimeric FoldOn of T4 fibritin, the Avitag sequence, and a hexahistidine affinity tag^23^, and cloned into the pCMV Sport6 expression vector. HA expression plasmids were co-transfected with a plasmid encoding cell surface expressed A/Puerto Rico/8/1934 neuraminidase (NA) into Expi293F cells (Thermo Fisher) with PEI Max reagent (Polysciences), following the manufacturer’s instructions. NA cleaves sialic acid and facilitates the release of the rHA into the supernatant, increasing protein yields. After 4 days, the supernatant was clarified by centrifugation. HA proteins were purified using Ni-NTA agarose resin (Qiagen) and gravity flow chromatography columns (Bio-Rad). Purified HA was buffer-exchanged and concentrated using centrifugal filter units with a 30 kDa molecular weight cutoff (Millipore). Purified HA proteins were stored at -80°C.

### Enzyme-linked immunosorbent assay (ELISA)

ELISA was performed on 96-well Immulon 4HBX extra high binding flat-bottom plates (Thermo Fisher). Plates were coated with phosphate-buffered saline (PBS) or recombinant HA protein from A/dairy cattle/Texas/24-008749-002-v/2024(H5N1) at 2 μg/mL in DPBS at 4 °C overnight. Plates were washed three times with 1X PBS+0.1% Tween-20 and then blocked for 1 h at room temperature (RT) using blocking buffer (1X TBS, 0.05% Tween-20, and 1% bovine serum albumin). Bovine serum samples were heat-inactivated at 56 °C for 1 h and then serially diluted fourfold in dilution buffer (1X TBS, 0.05% Tween-20, and 0.1% bovine serum albumin). Bovine milk samples were serially diluted two-fold in dilution buffer (1X TBS, 0.05% Tween-20, and 0.1% bovine serum albumin). After blocking, plates were washed three times with 1X PBS+0.1% Tween-20. Diluted milk samples were then transferred to ELISA plates and incubated for 2 h at RT. After washing three times with 1X PBS+0.1% Tween-20, peroxidase-conjugated goat anti-bovine IgG polyclonal antibody (Jackson ImmunoResearch Laboratories) was added to the corresponding wells at a concentration of 1:5000 in dilution buffer for 1 h at RT. After washing three times with 1X PBS+0.1% Tween-20, all plates were developed by adding SureBlue TMB peroxidase substrate (SeraCare) for 5 minutes at RT, followed by stopping the reaction with 250 mM HCl solution. The absorbance was measured at 450 nm using a SpectraMax ABS Plus plate reader (Molecular Devices). Background OD values from the plates coated with PBS were subtracted from the OD values from plates coated with recombinant HA protein. Endpoint titers were calculated as the reciprocal serum dilution that generated an equivalent optical density (OD) of 0.5.

### Foci Reduction Neutralization assay

Foci reduction neutralization (FRNT) assays were carried out using conditionally replicative influenza viruses that encode GFP instead of PB1, as previously described^24^. Conditionally replicative viruses were generated encoding the hemagglutinin (HA) and neuraminidase (NA) from A/dairy cattle/Texas/24-008749-002-v/2024(H5N1). Bovine serum samples were pretreated with receptor-destroying enzyme (RDE) (Denka Seiken) for 2 h at 37 °C, followed by heat inactivation at 56 °C for 30 min. RDE-treated sera were serially diluted in neutralization assay media (NAM, medium 199 supplemented with 0.01% heat-inactivated FBS, 0.3% BSA, 100 U of penicillin/mL, 100 µg of streptomycin/mL, 100 µg of calcium chloride/mL, and 25 mM HEPES), mixed with virus, and incubated at 37 °C for 1 h. After incubation, 2.5x10^4^ MDCK-SIAT1-CMV-PB1-TMPRSS2 cells were added to each well. Plates were incubated for 40 h at 37 °C followed by measurement of GFP fluorescence intensity on an EnVision multi-mode microplate reader (PerkinElmer) using an excitation wavelength of 485 nm and an emission wavelength of 515 nm. Neutralization titers were expressed as the reciprocal of the highest dilution of sera that inhibited 90% of GFP expression relative to control wells with no sera.

### Hemagglutination inhibition (HAI) assay

For HI assay, 50 µL of bovine serum was treated with 150 µL of RDE (Denka Seiken) with an overnight incubation at 37°C in a water bath, followed by 300 µL of saline solution (0.85%), and heat inactivation at 56°C for 30 minutes. RDE-treated serum samples were adsorbed with 100% rooster red blood cells for 1 hour at 4°C to remove nonspecific agglutinins. RDE-treated serum samples were run in HI assays using 0.5% rooster red blood ^25^. The HAI titers were reported as geometric mean titer.

### Mapping of antibody specificity by deep mutational scanning

Production of H5 HA deep mutational scanning libraries has been described previously^21^. The libraries are based on the HA of a 2.3.4.4b clade A/American Wigeon/South Carolina/USDA-000345-001/2021 strain, with mutants of this HA expressed on the surface of pseudotyped lentiviral particles (pseudoviruses). Importantly, these pseudoviruses can only undergo a single round of cell entry and are therefore not fully infectious pathogens capable of causing disease, thereby providing a safe way to study the impacts of HA mutations. To identify mutations that affect neutralization by the cattle sera deep mutational scanning libraries were incubated with serum concentrations that were sufficient to neutralize between 50% and 90% of the pseudovirus libraries, as estimated via neutralization standard used in deep mutational scanning experiments. After 45 min incubation at 37°C, the sera-virus mix was used to infect 293T cells. At 15 h after infection, non-integrated viral genomes were recovered and prepared for Illumina sequencing as described previously^21^. A non-neutralizable standard that was added to the libraries was used to determine effects of mutations on neutralization^21^. Neutralization escape for individual libraries was calculated using a biophysical model implemented in polyclonal package^26^. For each serum experiments were performed with two biological library replicates. Before use sera were treated with receptor destroying enzyme for 2 h at 37°C and heat-inactivated for 30 min at 56°C.

The data analysis pipeline for deep mutational scanning is available at https://github.com/dms-vep/Flu_H5N1_American-Wigeon_2021_HA_DMS_cattle_Penn_sera. Numerical values for individual sera neutralization are at https://github.com/dms-vep/Flu_H5N1_American-Wigeon_2021_HA_DMS_cattle_Penn_sera/blob/master/results/summaries/cattle_sera_escape_per_antibody_escape.csv. H3 numbering is used in this analysis, spreadsheet with conversion to different numbering schemes is at https://github.com/dms-vep/Flu_H5N1_American-Wigeon_2021_HA_DMS_cattle_Penn_sera/blob/master/data/site_numbering_map.csv.

To assess the conservation at key amino-acid sites targeted by the neutralizing activity of the sera, we downloaded all H5 HA proteins in GISAID as of Oct-27-2025 (see GISAID EPI_SET_251028bx, EPI_SET_251028no, and EPI_SET_251028mu for acknowledgments). We then calculated the frequency of each amino acid at each site among all H5 HAs, clade 2.3.4.4b HAs, and dairy cow HAs.

### Viral RNA detection in clinical samples

Viral RNA was extracted from nasal swabs, ocular swabs, urine, saliva, sera, and liquid collected from thawed diaphragm tissue using MagMAX™ CORE Nucleic Acid Purification Kit (Thermo Fisher Scientific, MA) with an extra wash using 80% ethanol. A piece of approximately 10 mm of mammary gland tissue from each quarter and supramammary lymph nodes were processed separately with the clarifying solution of the MagMAX™ CORE Mechanical Lysis module for tissue homogenization in GentleMACS™ Octo Dissociator (Miltenyi Biotec). Following tissue dissociation, viral RNA was extracted with the same protocol described above. Viral RNA was extracted from milk samples using MagMAX™ CORE Mastitis & Panbacteria Module (Thermo Fisher Scientific). Whole blood samples were processed for RNA extraction using LeukoLOCK™ Total RNA Isolation System (Thermo Fisher Scientific).

All RNA extracted were assessed via PCR for viral RNA detection using a commercial RT–qPCR VetMAX™-Gold SIV Detection Kit (Thermo Fisher Scientific). Milk samples that were PCR positive with Ct ≤ 35 were submitted to USDA-Animal and Plant Health Inspection Service (APHIS) National Veterinary Servies Laboratories (NVSL) lab to perform virus isolation in 10-day-old embryonating chicken eggs to determine the presence of viable virus. *Ct* values from RT-qPCR of 10-fold diluted stock virus of known TCID_50_ titer were extrapolated into a virus titration curve using GraphPad Prism (v.10.3.0) and reported as Log_10_ TCID_50_/mL equivalent Ct.

### Pathologic assessment of mammary gland tissues

At necropsy, mammary gland quarters were identified (front left, front right, rear left, rear right) by color markings, and the entire gland was dissected from the abdominal wall. For macroscopic evaluation, each quarter was dissected into cranial, middle, and caudal sections that were assigned a percentage of fibrosis score (percent of section that was fibrotic) which were then averaged by quarter. Out of each cranial, middle, and caudal section, three representative histologic sections were taken for a total of nine histologic sections per quarter and thirty-six sections per cow. Formalin-fixed tissues were transferred to 70% ethanol 48 h post-collection and processed routinely for microscopic evaluation. Tissues were stained by hematoxylin and eosin stain and Picrosirius Red stain per manufacturer’s instructions (AbCam) to highlight fibrosis^27^. Each section (36 per cow) was scored using a dual-component, intra- and interlobular fibrosis, histologic scoring rubric that was developed to assess mammary fibrosis. Each component was independently scored based on two criteria: histologic features (scale 0–4) and the percentage of tissue affected (scale 0–4). These scores were summed to produce a component-specific score ranging from 0 to 8. The overall fibrosis score was the combined total of both components (intra- and interlobular fibrosis), yielding a maximum possible score of 16. The results by quarter were summed for a possible total of 144 and then averaged by status (uninoculated or inoculated) of gland within a cow (Supplementary Table 6). Immunohistochemical staining (IHC) targeting the NP antigen (GeneTex) of IAV was performed on all sections that originated from an uninoculated quarter that had viral RNA detected at any timepoint, as previously described^28^.

### Statistical Analysis

Antibody data analyses were performed using GraphPad Prism Software Version 10 (GraphPad Software Inc.) using an unpaired *t*-test. Percent change in rumination, milk production, and milk PCR in the uninoculated quarters, were analyzed with two-way repeated measures ANOVA followed by Tukey’s multiple comparisons test to compare vaccinated versus unvaccinated groups across dpi. Viral RNA detection by RT-qPCR in the inoculated quarters were analyzed by a mixed-effects model (REML) followed by Tukey’s multiple comparisons test. Differences in mammary microscopic lesions (fibrosis) and mammary macroscopic lesions between vaccinated and unvaccinated cows were evaluated using the Mann–Whitney U test. Statistical significance was evaluated using an alpha threshold of 0.05 (*). *P* values greater than 0.05 were considered not significant (ns). Pearson’s correlation coefficients in unvaccinated and vaccinated lactating cows were calculated between Ct values (milk bucket, milk from inoculated and uninoculated quarters) and parameters: milk production (n = 56), rumination time (n = 58), CMT scores from inoculated quarters (n = 60) and uninoculated quarters (n = 60), milk color from inoculated quarters (n = 60) and uninoculated quarters (n = 60), and milk consistency scores from inoculated quarters (n = 60) and uninoculated quarters (n = 60), across all time points, using GraphPad Prism Software Version 10 (GraphPad Software Inc.). Correlations were considered significant at *P* < 0.05.

## Data Availability

Source data are provided with the manuscript or made publicly available.

## Code Availability

Code for data analysis has been made publicly available, and links for the source code have been provided with the manuscript.

## Acknowledgements

We thank the staff at the New Bolton Center at the University of Pennsylvania. In particular, we acknowledge the technical support provided by Katie Basiilio, Mark Roney, and Payne Vermillion at the New Bolton Center Marshak Dairy Herd, and New Bolton Center laboratory support from E. Yvonne Miller-Lux and Betty Osborne. We thank the USDA NADC and NVSL leadership and animal resource and facilities and engineering units. In particular, we thank Katharine Young, Emily Love and Sarah Anderson for laboratory technical assistance; Tonia McNunn for compliance; Denise Chapman, Michelle Crocheck, Jenny Rasmussen, Tiffany Williams, Jonathan Gardner, Jared Peterson, Emma Hay, Emma Pratt, as well as multiple personnel from NADC for their expertise, support and assistance with animal studies in the high containment BSL-3Ag facility. This work was supported in part by the National Institute of Allergy and Infectious Diseases, National Institutes of Health, Department of Health and Human Services (contract numbers 75N93021C00015), the USDA ARS (project number 5030-32000-231-000-D), and USDA APHIS (contract number 3200-231-112-I). D.G., G.Z and C.S. were supported by an appointment to the USDA-ARS Research Participation Program administered by the Oak Ridge Institute for Science and Education (ORISE) through an interagency agreement between the U.S. Department of Energy (DOE) and the USDA under contract no. DE-AC05-06OR23100. Mention of trade names or commercial products in this article is solely for the purpose of providing specific information and does not imply recommendation or endorsement by the US Government. USDA is an equal opportunity provider and employer.

## Supplementary files

**Supplementary Table 1**. Date of birth and age of lactating cows in the study.

**Supplementary Table 2**. Virus isolation from milk samples with qPCR Ct values < 35.5.

**Supplementary Table 3**. Macroscopic lesion score in uninoculated mammary quarters.

**Supplementary Table 4**. Microscopic lesion score of uninoculated mammary quarters of vaccinated and unvaccinated lactating cows.

**Supplementary Table 5**. Primers and probe sequences for A/dairy cattle/Texas/24-008749-002/2024(H5N1) HA (co-Tx/24 H5 HA) mRNA.

**Supplementary Table 6**. Microscopic lesion scoring system for fibrosis.

## References

1. Nguyen, T.Q. et al. Emergence and interstate spread of highly pathogenic avian influenza A(H5N1) in dairy cattle in the United States. Science 388, eadq0900 (2025).

2. Pena-Mosca, F. et al. The impact of highly pathogenic avian influenza H5N1 virus infection on dairy cows. Nat Commun 16, 6520 (2025).

3. Mostafa, A. et al. Avian influenza A (H5N1) virus in dairy cattle: origin, evolution, and cross-species transmission. mBio 15, e0254224 (2024).

4. Baker, A.L. et al. Dairy cows inoculated with highly pathogenic avian influenza virus H5N1. Nature 637, 913–920 (2025).

5. Halwe, N.J. et al. H5N1 clade 2.3.4.4b dynamics in experimentally infected calves and cows. Nature 637, 903–912 (2025).

6. Souza, C.K. et al. H5 influenza virus mRNA-lipid nanoparticle (LNP) vaccination elicits adaptive immune responses in Holstein calves. bioRxiv, 2025.2005.2001.651548 (2025).

7. Burrough, E.R. et al. Highly Pathogenic Avian Influenza A(H5N1) Clade 2.3.4.4b Virus Infection in Domestic Dairy Cattle and Cats, United States, 2024. Emerg Infect Dis 30, 1335–1343 (2024).

8. Caserta, L.C. et al. Spillover of highly pathogenic avian influenza H5N1 virus to dairy cattle. Nature 634, 669–676 (2024).

9. Song, H. et al. Receptor binding, structure, and tissue tropism of cattle-infecting H5N1 avian influenza virus hemagglutinin. Cell 188, 919–929 e919 (2025).

10. Nelli, R.K. et al. Exploring influenza A virus receptor distribution in the lactating mammary gland of domesticated livestock and in human breast tissue. J Dairy Sci (2025).

11. Center for Veterinary Biologics, USDA. Notification of U.S. Department of Agriculture’s (USDA) Request for Information for Highly Pathogenic Avian Influenza (HPAI) Vaccines for Use in Cattle, Notice No. 24-09. (2024).

12. Zhang, S., Liu, G., Guo, A. & Chen, Y. Maternal antibody transfer efficiency: The impact of M. bovis-BoHV-1 combined vaccine. Virology 611, 110656 (2025).

13. Abousenna, M.S., Shafik, N.G. & Abotaleb, M.M. Evaluation of humoral immune response and milk antibody transfer in calves and lactating cows vaccinated with inactivated H5 avian influenza vaccine. Sci Rep 15, 4637 (2025).

14. Wiggins, J., Madapong, A. & Weaver, E.A. Dual-Route H5N1 Vaccination Induces Systemic and Mucosal Immunity in Murine and Bovine Models. bioRxiv, 2025.2009.2021.677614 (2025).

15. Furey, C. et al. Development of a nucleoside-modified mRNA vaccine against clade 2.3.4.4b H5 highly pathogenic avian influenza virus. Nat Commun 15, 4350 (2024).

16. Low, J.M. et al. Codominant IgG and IgA expression with minimal vaccine mRNA in milk of BNT162b2 vaccinees. NPJ Vaccines 6, 105 (2021).

17. Hanna, N. et al. Detection of Messenger RNA COVID-19 Vaccines in Human Breast Milk. JAMA Pediatr 176, 1268–1270 (2022).

18. Hanna, N. et al. Biodistribution of mRNA COVID-19 vaccines in human breast milk. EBioMedicine 96, 104800 (2023).

19. Lang, Y. et al. Detection of antibodies against influenza A viruses in cattle. J Virol 99, e0213824 (2025).

20. Dadonaite, B. et al. A pseudovirus system enables deep mutational scanning of the full SARS-CoV-2 spike. Cell 186, 1263–1278 e1220 (2023).

21. Dadonaite, B. et al. Deep mutational scanning of H5 hemagglutinin to inform influenza virus surveillance. PLoS Biol 22, e3002916 (2024).

22. Shi, J. et al. H5N1 virus invades the mammary glands of dairy cattle through ‘mouth-to-teat’ transmission. Natl Sci Rev 12, nwaf262 (2025).

23. Whittle, J.R. et al. Flow cytometry reveals that H5N1 vaccination elicits cross-reactive stem-directed antibodies from multiple Ig heavy-chain lineages. J Virol 88, 4047–4057 (2014).

24. Doud, M.B., Hensley, S.E. & Bloom, J.D. Complete mapping of viral escape from neutralizing antibodies. PLoS Pathog 13, e1006271 (2017).

25. Kitikoon, P., Gauger, P.C. & Vincent, A.L. Hemagglutinin inhibition assay with swine sera. Methods Mol Biol 1161, 295–301 (2014).

26. Yu, T.C. et al. A biophysical model of viral escape from polyclonal antibodies. Virus Evol 8, veac110 (2022).

27. Farris, A.B. & Alpers, C.E. What is the best way to measure renal fibrosis?: A pathologist’s perspective. Kidney Int Suppl (2011) 4, 9–15 (2014).

28. Arruda, B. et al. Divergent Pathogenesis and Transmission of Highly Pathogenic Avian Influenza A(H5N1) in Swine. Emerg Infect Dis 30, 738–751 (2024).

