## Supplemental Figures for "Evaluation of an H5 influenza virus mRNA-lipid nanoparticle (LNP) vaccine in lactating dairy cows"

### Extended Data Figure 1

a

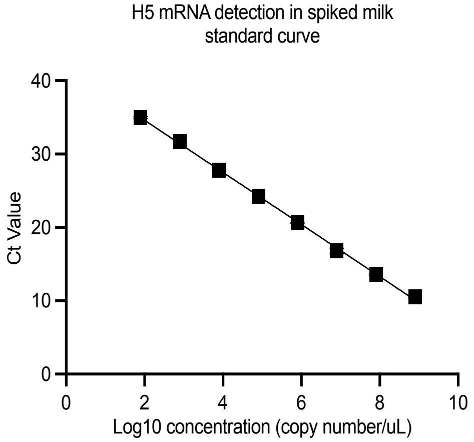


b

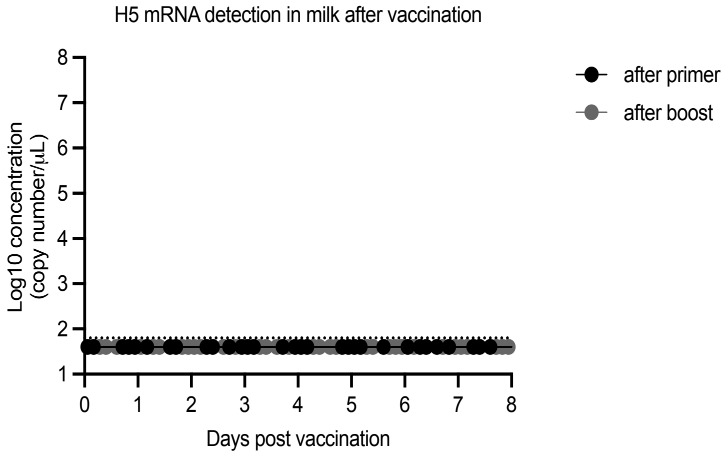

**Extended Data Figure 1. Absence of vaccine mRNA detection in milk after vaccination. (a)** standard curve of known concentration of mRNA spiked in milk and threshold cycle (Ct) measured using a real-time qPCR for H5 mRNA vaccine. **(b)** Real-time qPCR quantification of vaccine H5 mRNA from milk of vaccinated animals collected up to 8 days post-vaccination. Data shown are mean ± SEM of two replicates.

### Extended Data Figure 2

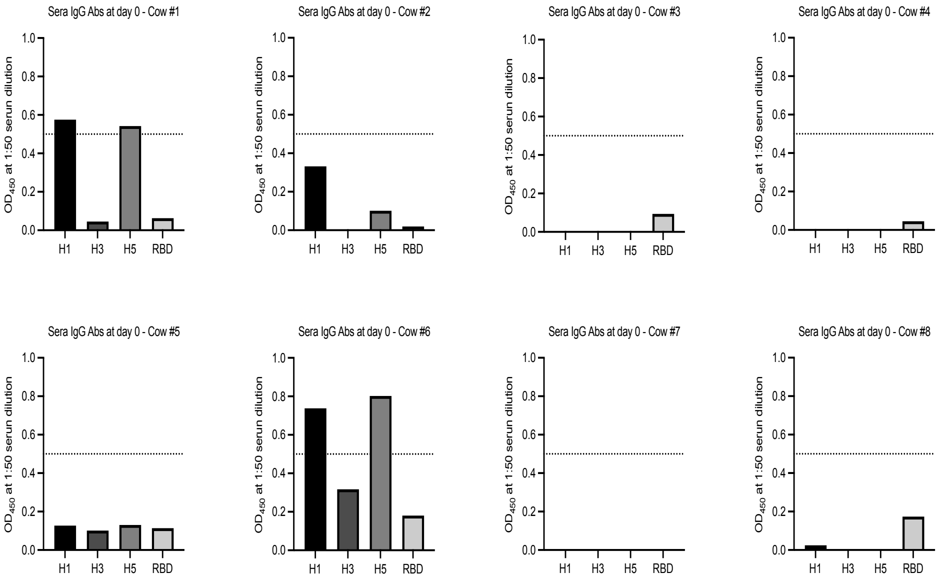

**Extended Data Figure 2. Evidence of influenza virus pre-exposure in sera from lactating dairy cows in the study.** Sera collected at day 0 from animals in the study were quantified by ELISA against influenza H1, H3, and H5 rHAs, as well as SARS-CoV-2 RBD for each animal.

### Extended Data Figure 3

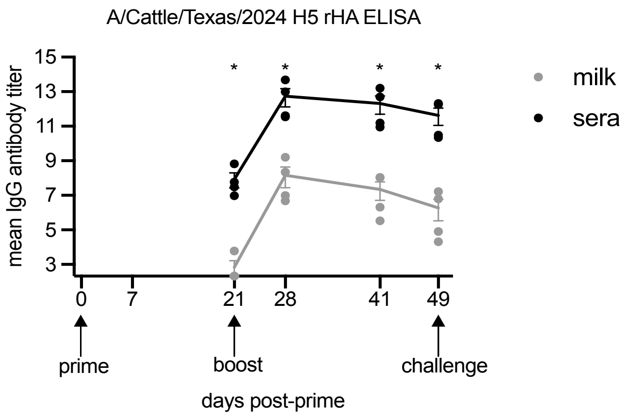

**Extended Data Figure 3. Comparison of binding antibodies in sera and milk of vaccinated animals.** Sera (closed black circles) or milk (closed gray circles) collected at days 21, 28, 41, and 49 from vaccinated cows were quantified by ELISA. Data are mean ± SEM, and individual data points (n=4) are shown. Significant statistical differences ($P$<0.05) between groups were determined using an unpaired *t*-test and are indicated by an asterisk.

### Extended Data Figure 4

| | | Milk PCR | | VI on milk | | Mammary tissue PCR | | Mammary tissue IHC | SMLN tissue PCR | SMLN tissue PCR | Diaphragm PCR | Other clinical samples - PCR | | | | | |
|---|---|---|---|---|---|---|---|---|---|---|---|---|---|---|---|---|---|
| **Cow** | **Group** | **inoculated** | **uninoculated** | **inoculated** | **uninoculated** | **inoculated** | **uninoculated** | **uninoculated** | **Right** | **Left** | **DL** | **NS** | **OS** | **Urine** | **Saliva** | **Sera** | **LL** |
| 1 | unvaccinated | positive | positive | positive | positive | postive | negative | negative | positive | positive | suspect | suspect | positive | negative | negative | suspect | suspect |
| 2 | unvaccinated | positive | negative | positive | negative | postive | negative | negative | positive | positive | negative | negative | positive | suspect | negative | suspect | negative |
| 3 | unvaccinated | positive | negative | positive | negative | postive | negative | negative | negative | negative | negative | negative | suspect | positive | negative | suspect | negative |
| 4 | unvaccinated | positive | positive | positive | positive | postive | positive | positive | negative | positive | negative | positive | negative | positive | suspect | suspect | negative |
| 5 | vaccinated | positive | negative | postive | negative | postive | negative | negative | positive | positive | negative | negative | negative | negative | negative | negative | negative |
| 6 | vaccinated | positive | negative | postive | negative | postive | negative | negative | positive | positive | negative | negative | negative | negative | negative | negative | positive |
| 7 | vaccinated | positive | negative | postive | negative | negative | negative | positive | negative | negative | negative | suspect | negative | negative | negative | negative | negative |
| 8 | vaccinated | positive | negative | postive | negative | negative | negative | negative | negative | negative | negative | negative | negative | negative | negative | negative | negative |

**Extended Data Figure 4. Evidence of virus migration from an inoculated to uninoculated mammary quarters.** Red represents positive, yellow suspect and white negative for each test, including qPCR on milk, virus isolation on milk; VI, mammary tissues, other clinical samples and IHC. SMLN, supramammary lymph node; DL, Diaphragm liquid; NS, nasal swab; OS, ocular swab; LL, LeucoLOCK from whole blood; NA refers to not applicable.

### Extended Data Figure 5

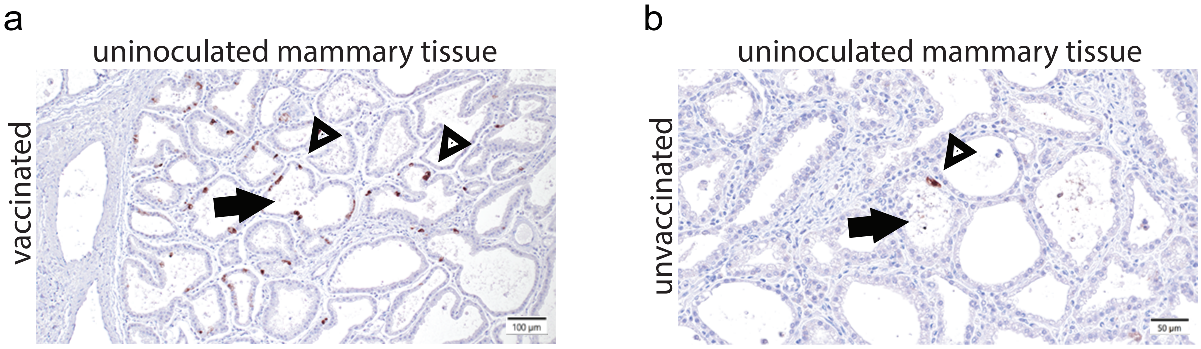

**Extended Data Figure 5.** Detection of highly pathogenic avian influenza virus by immunohistochemistry in uninoculated mammary quarters. Immunolabeling of numerous secretory epithelial cells (arrowhead) and cellular debris in the lumens of secretory alveoli (arrow) (a; Cow 7, vaccinated, rear right quarter, uninoculated). Immunolabeling of a secretory epithelial cell (arrowhead) and cellular debris in the lumen of a secretory alveoli (arrow) (b; Cow 4, front left quarter, unvaccinated uninoculated).

### Extended Data Figure 6

a

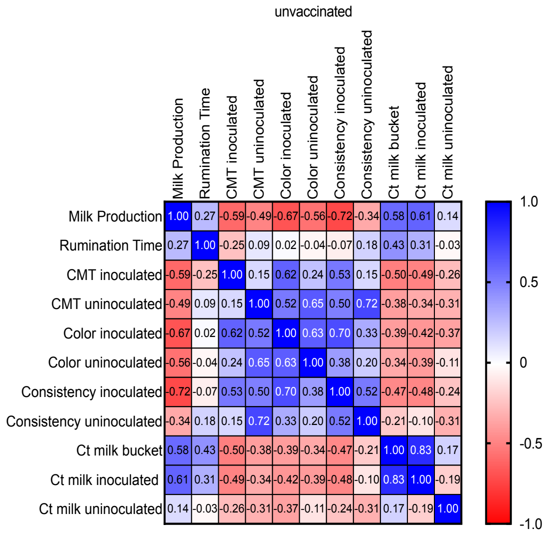


b

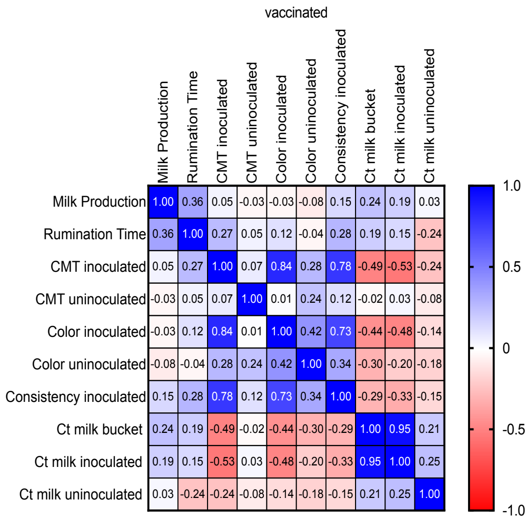

**Extended Data Figure 6. Correlation between virus RNA detection and clinical signs in unvaccinated and vaccinated lactating cows.** Black open boxes highlighted the significant (*P*<0.05, two-sided) Pearson's correlation coefficients in unvaccinated **(a)** and vaccinated **(b)** lactating cows are shown between Ct values (milk bucket, milk from inoculated and uninoculated quarters) and milk production , rumination time , CMT scores from inoculated and uninoculated quarters , milk color from inoculated and uninoculated quarters , and milk consistency scores from inoculated and uninoculated quarters , across all time points.
